# FM3VCF: A Software Library for Accelerating the Loading of Large VCF Files in Genotype Data Analyses

**DOI:** 10.1101/2023.06.25.546413

**Authors:** Zhen Zuo, Qi Li, Zhuo Li, You Tang, Meng Huang

**Author notes:** Corresponding author E_mail (TY).

## Abstract

**Background:** The increasing size of genotype data has led to the loading of VCF files becoming a computational bottleneck in various analyses, including imputation and genome-wide association studies (GWAS). To address this issue, we developed a software library, FM3VCF (fast M3VCF), that utilizes multiple CPU threads to accelerate this process.

**Findings:** FM3VCF can convert VCF files into the exclusive data format of MINIMAC4[1], M3VCF[1], and efficiently read and parse data from VCF files. In comparison to m3vcftools[1], FM3VCF is approximately 20 times faster for compressing VCF files to M3VCF format. Furthermore, FM3VCF is approximately 3 times faster than HTSlib[2], including decompressing and parsing, for reading compressed VCF files. FM3VCF is written in C and is open-source, available for download from https://github.com/Oliver-111/m3vcf under the MIT/BSD license.

**Conclusion:** FM3VCF is a powerful tool for accelerating the loading of large VCF files in genotype data analyses. By fully utilizing multiple CPU threads, FM3VCF can significantly reduce the computational burden in various genomic analyses.

## Background

The Variant Call Format (VCF)[3] has emerged as a widely used format for storing variation data, thanks to its flexibility and unambiguous nature that allows for additional information to be included. However, with the advent of large-scale projects involving thousands of samples and millions of variants, the size of VCF files has grown significantly, even in compressed format. This has led to higher costs associated with loading and compression of such files. To tackle this issue, the M3VCF format was developed, which specifically aims to reduce the size of VCF files. Currently, M3VCF is the only format supported by the reference panel in MINIMAC4[1], a software tool designed for efficient imputation of large-scale sequencing data. Recently, Zhang et al. (2021) conducted a study to compare the compression performance of m3vcftools with that of other commonly used compression methods, such as M3VCF, GZIP[4], BZIP2[5], and LZMA[6]. The results showed that m3vcftools achieved the highest compression ratios across all datasets tested, including those with large numbers of samples and variants. Furthermore, m3vcftools outperformed other compression methods in terms of compression and decompression speed.

Large-scale studies such as Genome-Wide Association Studies (GWAS) and genotype imputation require efficient access to variants from VCF or M3VCF files. While the m3vcftools is faster than MINIMAC3[1] for compressing VCF files to M3VCF format, it may not be suitable for very large datasets. Although the HTSlib library provides a high-performance interface in C language for accessing VCF files with options to manipulate data at the variant and sample levels, a faster library is required for handling high-density markers, such as whole-genome sequencing data, in large-scale studies.

### Approach

We aimed to create a high-performance library, FM3VCF, for effectively compressing VCF files to M3VCF files by loading, parsing, and compressing the data. The compression task using m3vcftools involves three main steps: reading and parsing the VCF file data, compressing and converting the VCF file records to M3VCF file records, and writing the resulting data into the M3VCF file. As m3vcftools is single-threaded, all three compression tasks are completed within a single CPU thread. To improve performance, FM3VCF separates the Read, Compress, and Write processes and assigns them to different threads, enabling the three compression steps to be completed in parallel across multiple CPU threads. For example, if three threads could be used to read and compress three records. Compared to m3vcftools, which requires 9 steps, FM3VCF can complete the same three compression tasks in only 5 steps (Figure 1), leading to a significant increase in speed, especially for large data sets. The Compress step takes up the most time among the three processes, so a multi-threaded approach using pthread library[7] is adopted to run the Compress step in parallel across multiple CPU cores, resulting in more even distribution of processing time across the Read, Compress, and Write steps. Additionally, the Read step is divided into Reading Data and Parsing, with Parsing taking up the majority of the time. To accelerate this process, OpenMP library[8] is used to run Parsing in parallel across multiple threads (Figure 1), leading to a significant improvement in processing speed for the Reading step.

**Figure1.**
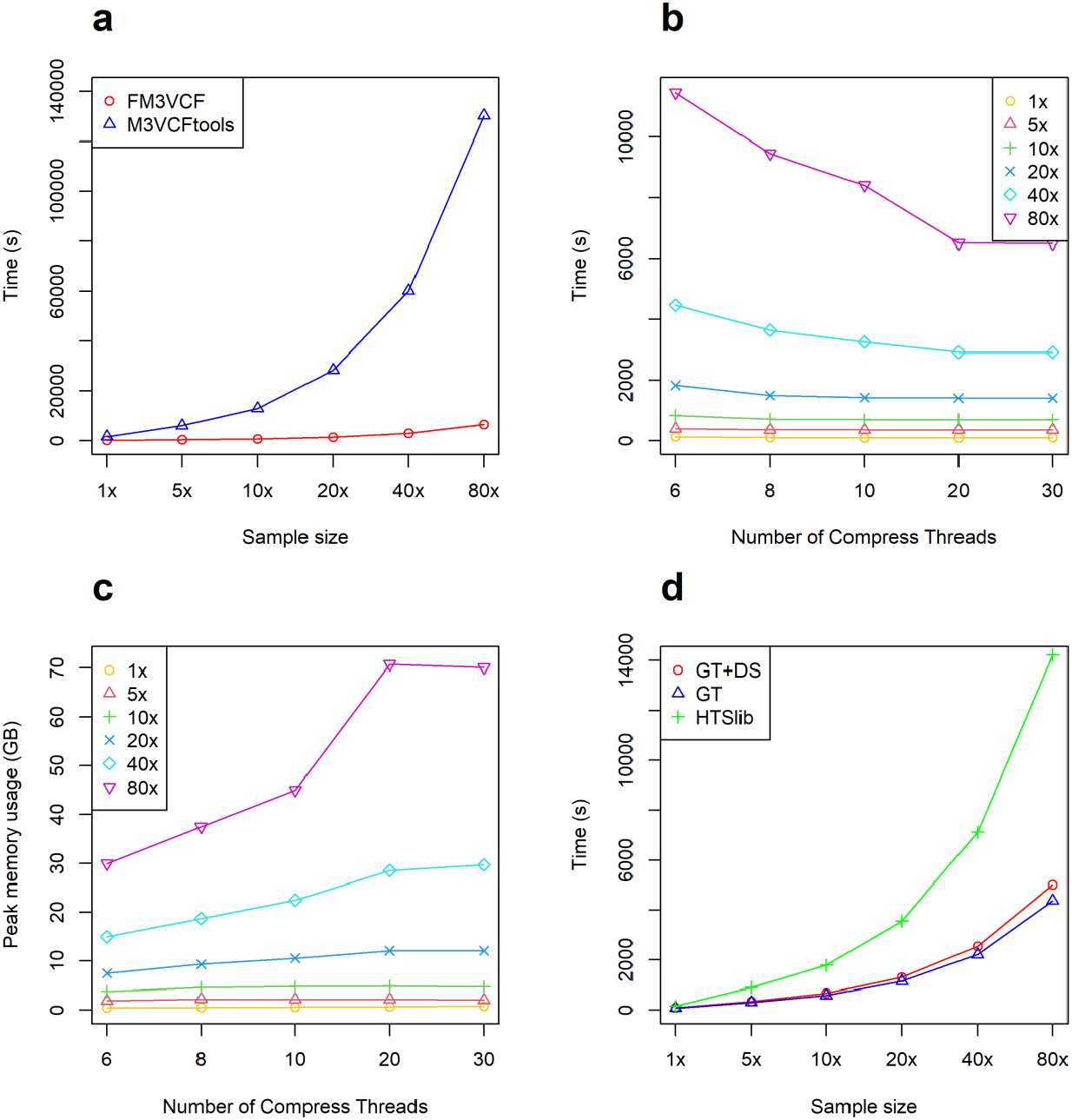
Performance evaluation of FM3VCF. (a) is the compression time for VCF files of different sizes is depicted in the figure. The blue line represents the results obtained using m3vcftools, while the red line represents the results obtained using FM3VCF. (b) illustrates the multithread computation time by FM3VCF. It includes results for sample sizes of 80x, 40x, 20x, 10x, 5x, and 1x, represented by the purple, cyan, blue, green, red, and orange lines, respectively. (c) showcases the multithread peak memory usage by FM3VCF. It presents the results for sample sizes of 80x, 40x, 20x, 10x, 5x, and 1x, denoted by the purple, cyan, blue, green, red, and orange lines, respectively.(d) displays the reading and parsing time for VCF files of different sizes. The green line corresponds to the reading time for different sample sizes using HTSlib. The red line represents the time required for parsing GT and DS by FM3VCF, while the blue line shows the time needed for parsing GT by FM3VCF.

### Usage

To illustrate the performance of M3VCF compression and VCF loading, we present examples of command line and library interface usage. It is important to note that M3VCF only accepts bi-allelic SNPs for compression. Detailed instructions, executable files, and source code can be accessed at https://github.com/Oliver-111/m3vcf.

VCF Reading and M3VCF Compression: To load a large number of genotype records from a VCF file using the library interface, the following steps are performed (Box 1): First, the VCF format structure is initialized (lines 4-6), followed by opening the VCF file (lines 7-9) and reading the header (line 10). There are two optional functions available for reading the variant records either variant by variant (line 11) or block by block (line 13), which allows efficient loading of large datasets. Finally, memory is released (line 18 or line 19), and the VCF file is closed (line 20). To specify the VCF file format, the parameter “FILE_MODE_GZ” is used for compressed files with the “.vcf.gz” suffix, while “FILE_MODE_NORMAL” is used for uncompressed files with the “.vcf” suffix in the vcfFileOpen() function. The VCF reading interface parses only the genotype (GT) and dosage (DS) information in the FORMAT field. Users can choose to load both by specifying the parameter “P_DS|P_GT” or load only GT by using “P_GT” in the vcfFileOpen() function. In the block reading function vcfFileReadDataBlock(), the number of variants in each block can be specified by the parameter “int numLines,” which enables the utilization of multiple threads for parsing.

After loading data from the VCF file, compression can be implemented using the following examples in Box 2, demonstrating the command line usage for compression (line 1) and decompression (line 2). The “-o” flag specifies the output file name, “compress” refers to M3VCF file compression, “-m/M” flag specifies whether the output file is compressed or not, and “convert” is used for decompressing M3VCF files to VCF format. For the library interface usage, the initial steps involve defining the block size (number of variants; line 4) and specifying the input and output file paths (lines 5-7). Then, the variable configuration for compression (lines 9-16) and decompression (lines 20-24) should be specified before launching the compression (lines 17-19) and decompression (lines 25-27) functions. After implementation, FM3VCF library automatically releases the allocated memory. For both compression and decompression variable configurations, the VCF (line 11, 24) and M3VCF file (line 13, 22) compression status should be specified, controlled by “FILE_MODE_NORMAL” and “FILE_MODE_GZ”. The compression function supports multiple threads, and users can specify the block size, number of threads, and memory size (lines 14-16) where the value 0 represents the optimized default setting (lines 15-16). Instructions for code compilation for Box 1-2 can be found at https://github.com/Oliver-111/m3vcf/blob/main/FM3VCF_instruction.pdf.

#### Box 1.

VCF reading and parsing (library interface)

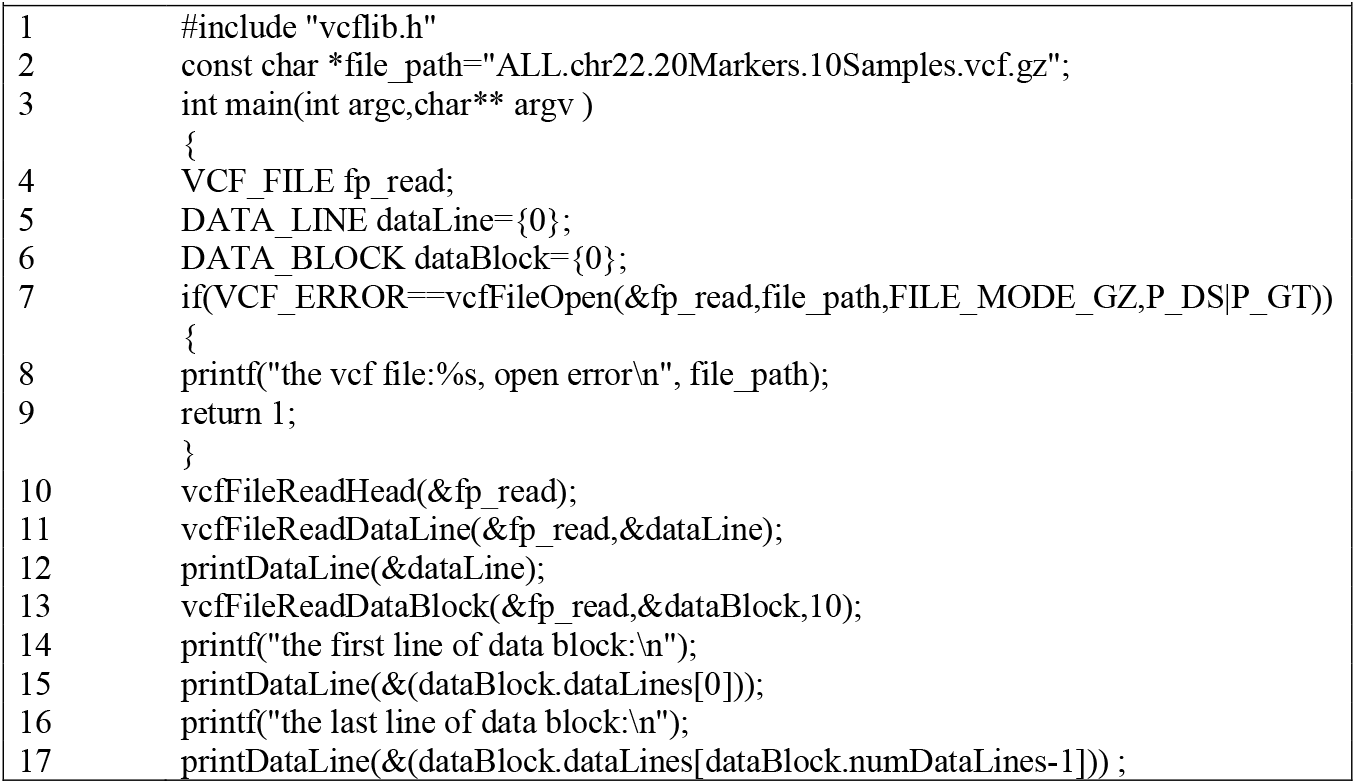
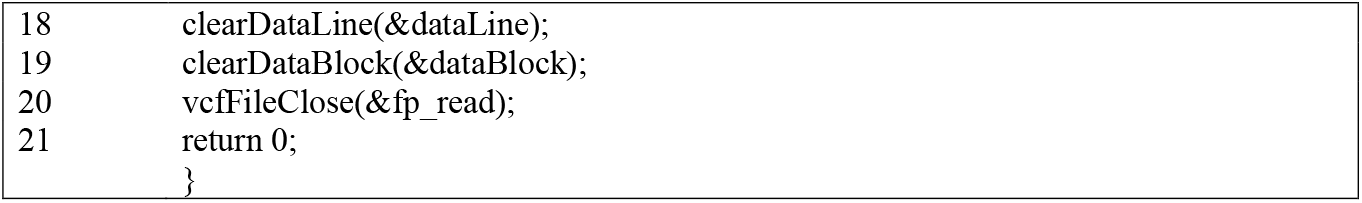

#### Box 2.

M3VCF compression (command line and library interface)

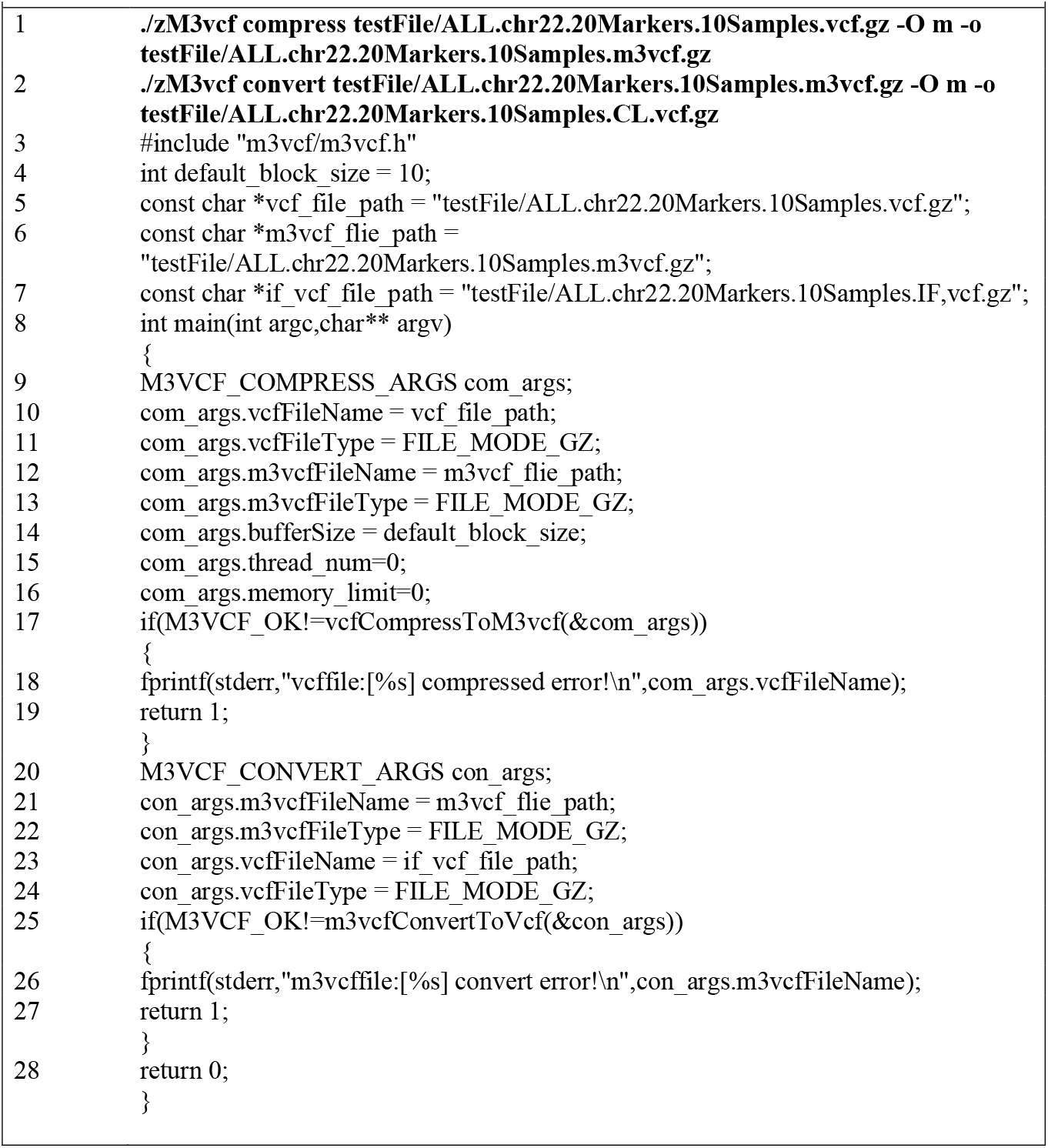

### Performance

In order to evaluate the performance of FM3VCF, we utilized a synthetic dataset consisting of 1,092 samples and 494,328 variants from chromosome 22 of the 1000 Genomes Project (http://ftp.1000genomes.ebi.ac.uk/vol1/ftp/release/20110521/ALL.chr22.phase1_release_v3.201_01123.snps_indels_svs.genotypes.vcf.gz; [9]) for speed testing. Five datasets were created by duplicating the original dataset with 5-, 10-, 20-, 40-, and 80-fold duplicated samples, respectively, for time-consuming tests.

For the compression of M3VCF files from compressed VCF files, we employed different numbers of threads (6, 8, 10, 20, and 30) to conduct the tests. For the dataset with 87,360 samples, m3vcftools took 36.1 hours for compression, while FM3VCF completed the task in 1.8 hours using 20 threads. Therefore, FM3VCF demonstrated approximately 20 times faster performance compared to m3vcftools (Figure 1a). With the number of threads increasing, the results indicated that using 20 threads reached saturation (Figure 1b), and the more threads, the more memory required (Figure 1c).

We also evaluated the performance of VCF file loading using the same datasets. When employing the block loading method, FM3VCF took 73 minutes, which was approximately 3 times faster than HTSlib with 87,360 samples and 10 threads, specifically for loading data and parsing GT only (Figure 1d). However, if both GT and DS were parsed, the time consumption remained almost the same as when parsing GT only (Figure 1d).

## Discussion

We have developed FM3VCF, a powerful library that allows efficient loading of data from VCF files and compression into M3VCF format. This software provides both a command line interface and a C/C++ library interface, offering researchers the flexibility and effectiveness needed to conduct genomic analysis using large datasets. With FM3VCF, researchers can effectively handle and analyze genomic data, enabling them to delve into complex genetic studies and derive valuable insights.

## Software Availability

The source code, instruction and usage demo could be found at: http://github.com/Oliver-111/m3vcf.

